# Drosophila blood cells bridge distant injury and gut homeostasis through Upd3-mediated inter-organ signaling

**DOI:** 10.1101/2025.10.13.681995

**Authors:** Yash Suresh Sheregare, Sandhya S. Visweswariah, Sveta Chakrabarti

## Abstract

Inter-organ communication is a central feature of host defence and tissue repair. In *Drosophila*, hemocytes secrete the cytokine-like ligand Unpaired 3 (Upd3, an IL-6 homolog) to activate JAK/STAT signaling in the gut, yet why this axis is essential for survival after injury has remained unclear. Here we show that loss of hemocyte activation by reactive oxygen species (ROS) and the consequent failure to produce Upd3 leads to adherens junction disruption, intestinal barrier dysfunction, and increased lethality following clean injury. Hemocyte-derived Upd3 drives sustained STAT activation in the gut epithelium, promoting enterocyte turnover and survival after injury. Chronic STAT activity further modulates intestinal stem cell fate and differentiation, indicating that hemocyte–gut signaling shapes long-term epithelial homeostasis. Notably, hemocytes home to the gut after distant wounding, where their localized presence enhances resistance to enteric infection. Taken together, these findings reveal that Drosophila blood cells bridge distant injury and gut homeostasis through Upd3-mediated inter-organ signaling that links wound sensing to epithelial integrity and host survival.

## Introduction

All animals comprise interconnected organ systems, and a perturbation to one organ impacts the functioning of another, thereby altering the physiology of the entire organism. Following external wounding, inter-organ communication between local wound sites and remote organs is crucial for host protection from the local wound (Lee & Miura, 2014). In Drosophila, a systemic wound response depends on the activation of circulating blood cells, or hemocytes, which secrete the IL-6–like cytokine Unpaired 3 (Upd3). Upd3 acts on multiple organs, including the gut, by activating the JAK/STAT signaling pathway (Agaisse et al., 2003; Chakrabarti et al., 2016; Yang et al., 2015). Because *Drosophila* lacks a closed vasculature, Upd3 released from hemocytes after injury diffuses through the open circulatory system to reach multiple tissues. However, which target organs are essential for survival following injury—and how they interpret this systemic cytokine signal—remains unknown.

We showed previously that intestinal epithelium renewal is a critical event for survival from septic injury (Chakrabarti et al., 2016). As the intestine in most organisms is in contact with the external environment, commensals, potential pathogens, and toxins, there must be a precise control of epithelial turnover to preserve barrier integrity, which is vital for organismal survival. The *Drosophila* intestine shares many structural and functional similarities with the mammalian small intestine and provides an accessible system to study epithelial homeostasis. Its epithelium is maintained by highly proliferative intestinal stem cells (ISCs) that give rise to progenitor enteroblasts (EBs), which in turn differentiate into absorptive enterocytes (Miguel-Aliaga et al., 2018; Trubin et al., 2024). Numerous studies have demonstrated that local and paracrine signals regulate ISC proliferation and differentiation in response to enterocyte loss or damage (Amcheslavsky et al., 2009; Biteau et al., 2008; Jiang et al., 2009; Medina et al., 2022; Shaw et al., 2010). However, whether distant cues—such as those from activated hemocytes—also contribute to intestinal repair and homeostasis remains poorly understood.

In mammals, nearly 70% of immune cells reside within or adjacent to the intestinal mucosa, highlighting the gut as a primary site of immune–epithelial communication and physiological regulation (Mowat & Agace, 2014; Vighi et al., 2008). Immune cells are typically thought to be mediators of pathogen defence, they may be key in other roles in tissue maintenance, driving mitotic activity and regeneration of intestinal epithelia (Arinda et al., 2022). In *Drosophila*, circulating hemocytes— functional analogs of mammalian macrophages—stimulate intestinal stem cell (ISC) proliferation following local epithelial damage (Ayyaz et al., 2015). Strikingly, even a wound at a distant site triggers ISC proliferation through the JAK/STAT-activating cytokine Unpaired 3 (Upd3) secreted from hemocytes (Chakrabarti et al., 2016). Thus, immune cells defend against infection and serve as systemic coordinators that couple tissue injury to regenerative programs in the intestine.

Efficient intestinal stem cell (ISC) proliferation is essential for survival after injury because sustained epithelial turnover preserves barrier integrity. When renewal is impaired, the gut barrier becomes compromised, allowing luminal microbes and toxins to breach the epithelium—a failure that is often lethal across metazoans (Chelakkot et al., 2018; Luissint et al., 2016; Marchiando et al., 2010; Panwar et al., 2021). In Drosophila, proper intestinal epithelial homeostasis relies on the continuous replacement of damaged or dying cells through apoptosis or apical extrusion (Amcheslavsky et al., 2009; Jiang et al., 2009; O’Brien et al., 2011; Zhai et al., 2018). Interestingly, a distant wound from the gut in flies induces caspase activation in enterocytes, leading to their removal and subsequent epithelial renewal (Amcheslavsky et al., 2020; Takeishi et al., 2013). These findings indicate that systemic signals can couple tissue injury to regenerative programs in the intestine, yet the source and identity of these long-range cues remain unclear.

Here, we investigate the role of hemocytes in coordinating intestinal repair following injury and examine how JAK/STAT activation in the gut contributes to organismal survival. We show that hemocyte-derived signals, initiated by reactive oxygen species (ROS) at the wound site, regulate ISC proliferation and epithelial renewal through the cytokine Unpaired 3 (Upd3). This inter-organ communication between immune and epithelial tissues establishes an equilibrium between proliferation, apoptosis, and survival, restoring gut homeostasis after injury. Furthermore, we demonstrate hemocytes home to the gut during recovery, where their presence enhances resistance to subsequent enteric infection. Collectively, our findings demonstrate that Drosophila hemocytes connect remote injury to gut homeostasis via Upd3-mediated inter-organ signaling, which associates wound detection with epithelial integrity and host survival.

## Results

### Adherens junctions are compromised upon injury in the absence of activated hemocytes

We recently demonstrated that hemocytes sense reactive oxygen species (ROS) at sites of injury, leading to the production of *upd3* and the release of Upd3, which in turn activates JAK/STAT signalling in the gut (Fig. S1A) (Chakrabarti & Visweswariah, 2020). This hemocyte-to-gut communication is critical for survival after sterile injury. Mutants defective in hemocyte activation, such as *Drpr^Δ5^* flies, fail to induce *upd3* expression and consequently lack Upd3 production, resulting in lethality upon injury. Significantly, this lethality cannot be attributed to septic infection, as immunodeficient flies (e.g., IMD pathway mutants) succumb more rapidly to bacterial challenge (Fig. S1B). These findings indicate that the Draper-dependent hemocyte response, acting through Upd3, is necessary to support gut recovery after injury.

Hemocytes secrete *upd3* that stimulates intestinal stem cell proliferation following a distant injury, even without infection (Chakrabarti et al., 2016). Since intestinal permeability is a marker of physiological age, we tested whether the absence of activated hemocytes in the fly is correlated with gut dysfunction (Rera, Azizi, et al., 2013; Rera et al., 2012; Rera, Clark, et al., 2013). We employed the Smurf assay, in which oral ingestion of blue dye reveals systemic leakage from the gut (Rera et al., 2012). Knockdown of *upd3* in hemocytes or enterocytes was tested (Fig. 1A, S1C). The efficacity of the knockdown was shown by our previous study and others (Agaisse et al., 2003; Chakrabarti et al., 2016; Yang et al., 2015). A progressive increase in Smurf-positive flies was observed only when Upd3 was depleted in hemocytes, but not in enterocytes following injury. Consistently, the *hmlΔGal4 > UAS-upd3-*RNAi flies displayed a higher proportion of Smurf-positive individuals than controls (Fig. 1A), indicating that hemocyte-derived Upd3 is essential to maintain gut barrier integrity after injury. This was not observed in the absence of any wound to the flies.

**Fig 1.**
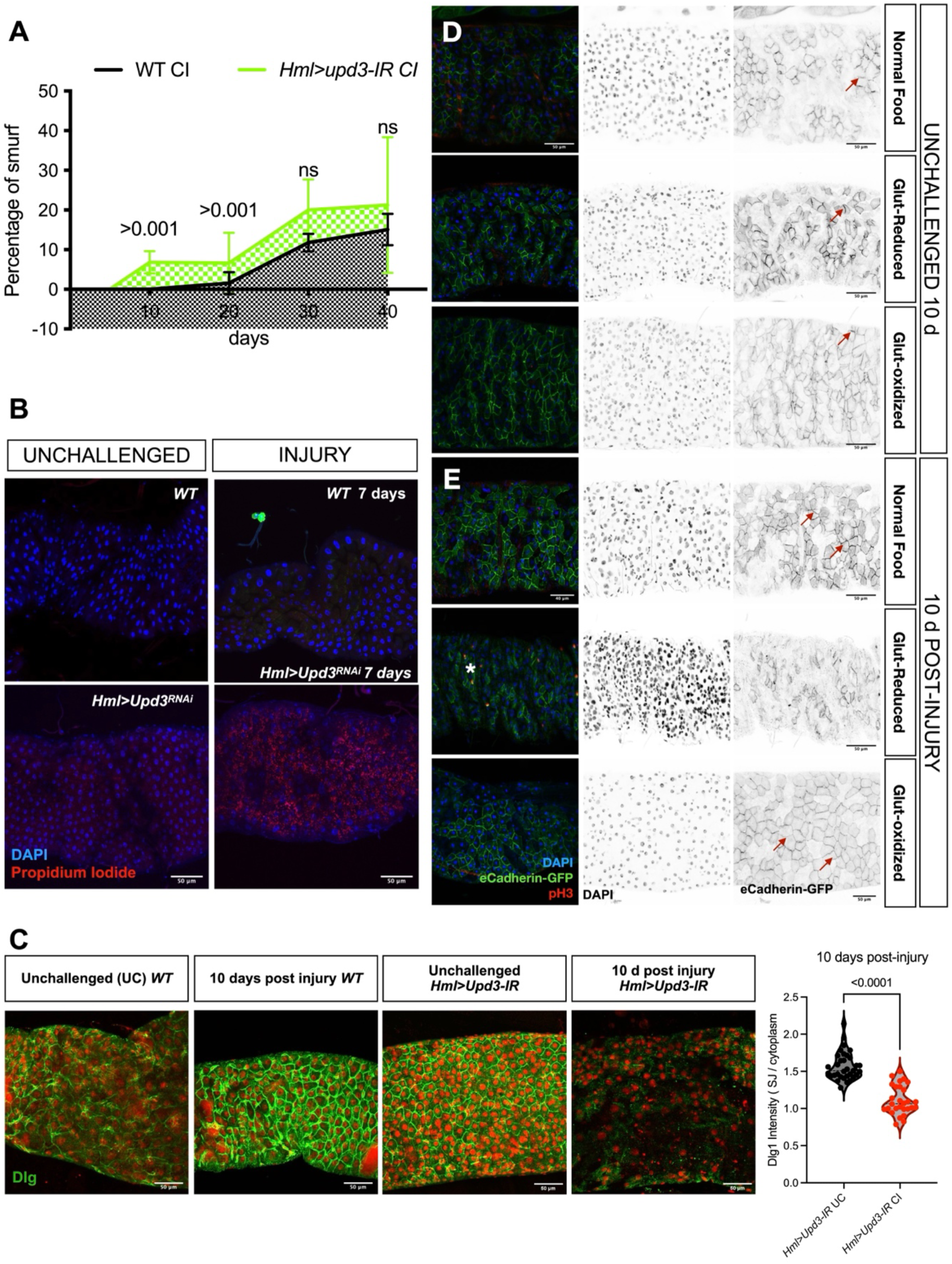
Hemocyte-derived *upd3* is required for intestinal barrier integrity. (A) Smurf assay of adult female flies maintained on food containing Blue dye no. 1. Flies with hemocyte-specific knockdown of *upd3* (*hmlΔ-GAL4* > *UAS-upd3-IR*; green curve) displayed an increased proportion of Smurf-positive individuals following clean injury (CI) compared to control flies (*hmlΔ-GAL4* > *w^1118^*; grey curve). (B) Propidium iodide (PI) feeding assay (see Methods). PI (red) entered enterocytes of *hmlΔ-GAL4* > *UAS-upd3-IR* flies 7 days post-injury, consistent with barrier leakage. In control guts, PI signal was largely excluded from enterocytes. Nuclei are counterstained with DAPI (blue). (C) Midgut epithelium from control (*hmlΔ-GAL4* > *w^1118^*) and *hmlΔ-GAL4* > *UAS-upd3-IR* flies stained for Discs large (Dlg, green) and DAPI (red). Disruption of cell junctions was observed in *upd3*-RNAi guts 10 days post-injury. Quantification of Dlg enrichment at bicellular junctions relative to cytoplasmic intensity revealed a significant reduction in junctional Dlg in *upd3*-RNAi midguts compared to controls (n = 31 UC, 30 CI; mean ± SD; P values shown are determined by Student’s t-test). (D) Localization of E-Cadherin in enterocytes of uninjured flies carrying an E-Cadherin-GFP reporter (E-Cad-GFP, green). Junctional localization was maintained before and after 10 days (red arrows). There was no observable difference between flies pre-fed with glutathione – oxidized and reduced form or normal food. Nuclei are counterstained with DAPI (blue), and mitotic cells are marked by anti-pH3 staining (red). (E) E-Cadherin localization in flies pre-fed with glutathione – oxidized and reduced form, for 1 day before injury. E-Cad-GFP showed mislocalization and internalization 10 days post-injury (white asterisk). Right panels show individual grayscale channels for E-Cad-GFP (green) and DAPI (blue). Scale bars, 50 μm. CI, clean injury; DAPI, 4′,6-diamidino-2-phenylindole; Dlg, Discs large; EC, enterocyte; GFP, green fluorescent protein; hml, hemolectin; IR, inverted repeat (RNAi construct); PI, propidium iodide; pH3, phospho-histone H3; RNAi, RNA interference; UAS, upstream activation sequence; WT, wild type.

To further assess epithelial barrier permeability, we adapted propidium iodide (PI) feeding as a tracer. PI is a small molecule (∼668 Da) widely used as a cell viability dye because it normally does not cross intact plasma membranes. Unexpectedly, when we administered PI to flies after thoracic wounding, it remained excluded from enterocytes in wild-type or uninjured flies but leaked into the cytoplasm of enterocytes in *Drpr^Δ5^* mutants and in flies lacking hemocyte-derived Upd3 (Fig. 1B, S2A, S2C). Importantly, we did not observe nuclear staining, consistent with PI reporting barrier leakage rather than cell death. Quantification confirmed a significant increase in PI uptake in *hmlΔGal4 > UAS-upd3-*RNAi guts relative to controls (Fig. S2A). These findings establish PI as a sensitive small-molecule tracer of gut barrier dysfunction in our system, complementary to larger molecular weight tracers such as 70 kDa FITC–dextran, which is excluded by the peritrophic matrix under normal conditions (Kuraishi et al., 2011).

We used another molecular marker, disc large 1 (dlg1), to monitor the junctional complex near the midgut cell apical pole (Fig. 1C). We silenced the *upd3* gene in hemocytes and monitored the localization of dlg1 in enterocytes from intestines 10 days post-injury. Enterocytes of flies where *upd3* was knocked down from hemocytes showed reduced levels of dlg1 at the junctions of enterocytes ten days post-injury compared to wild-type counterparts (Fig. 1C). Quantitative analysis of dlg1 fluorescence enrichment at bicellular junctions relative to cytoplasmic intensity confirmed a significant reduction in junctional dlg1 signal in *upd3*-RNAi midguts (Fig. 1C). Similar results were seen for the *Drpr^Δ5^* mutant flies (Fig. S2B). In addition, we checked Snakeskin (Ssk), a membrane protein associated with smooth septate junctions in the intestine. The localization of Ssk was disrupted in enterocytes of animals where *upd3* was knocked down from hemocytes 10 days post-injury (Fig. S2D).

We observed the localization of a gut epithelial junction protein, E-Cadherin, to gain insight into the molecular changes contributing to the increased barrier dysfunction rate in Upd3 knockdown flies. Because generating flies carrying both *E-Cad::GFP* and hemocyte-specific *upd3* RNAi was technically challenging, we established an alternative approach using glutathione feeding to modulate hemocyte *upd3* expression. We saw that flies fail to produce *upd3* ligand from their hemocytes post-injury if they are fed glutathione for a day before the injury (Figure S2E). Flies reared on normal food or food supplemented with oxidized glutathione showed membrane localization of E-Cadherin-GFP (E-Cad::GFP) in enterocytes before and ten days after an injury (Fig. 1D, E). In contrast, food combined with the antioxidant glutathione one day before injury led to E-Cad::GFP internalization and mislocalization from the cell membrane of enterocytes 10 days after the injury (Fig. 1D, E).

Taken together, the loss of Ssk, dlg1, and cadherin (E-Cad) in the absence of Upd3 produced from hemocytes following distal injury may lead to cellular permeability defects, which could hasten the loss of gut barrier function due to decreased cell integrity.

### Distant injury triggers intestinal stem cell proliferation

Recent work has shown intestinal epithelia maintain tissue integrity through dynamic cellular renewal and that disruption of this balance contributes to age-related barrier decline. An “age mosaic” of enterocytes, generated through continuous stem-cell–driven turnover, can delay epithelial ageing and preserve barrier function (Qin et al., 2025). However, excessive or uncontrolled intestinal stem cell (ISC) proliferation leads to epithelial hyperplasia and compromised barrier integrity (Buchon et al., 2009).

In Drosophila, injury triggers ISC proliferation through the cytokine Upd3, but whether this response depends on microbial cues remained unclear. Because changes in microbiota composition during ageing have been linked to ISC hyperactivation (Buchon et al., 2009), we asked whether the injury-induced proliferative response required microbial presence. In axenically reared wild-type flies, we observed no difference in the number of mitotic ISCs compared to conventionally reared flies following clean injury (Fig. 2A). However, in axenic *upd2,3^Δ^* mutant flies, mitotic ISCs were substantially reduced upon clean injury (Fig. 2A). Hence, Upd3 activates ISC proliferation independent of the presence of microbiota, underscoring its role as a direct inter-organ signal linking wounding to gut regeneration.

**Fig 2.**
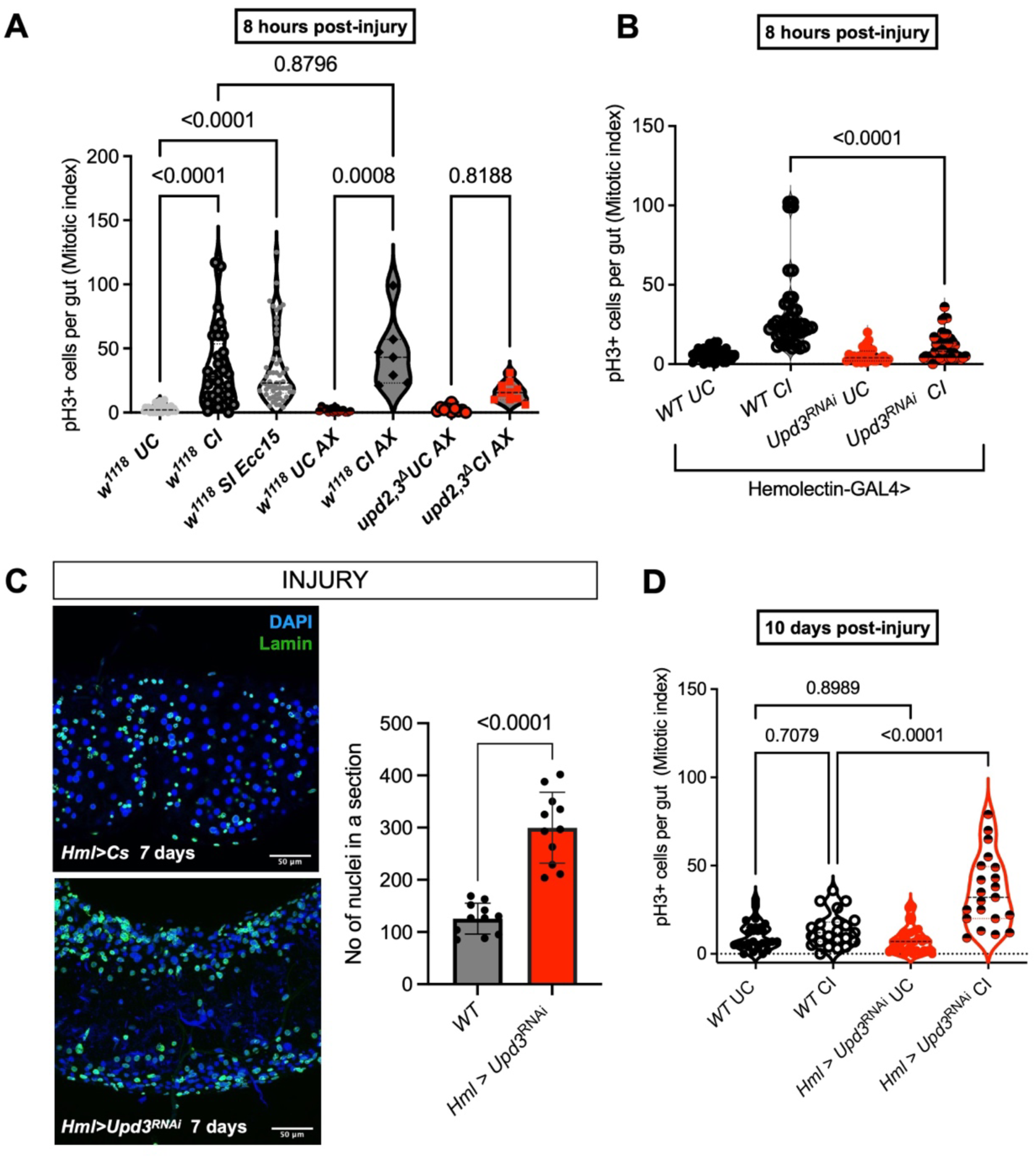
Injury activates stem-cell proliferation in the *Drosophila* intestine. (A) The mitotic index (as measured by Phospho-Histone-3-positive cells (pH3+) of the midgut wild-type female flies (*w^1118^*) 8 h post-injury and septic injury with *Erwinia caratovora caratovora 15* (*Ecc15*) displays increased proliferation as compared to the wild type (*w^1118^*) unchallenged counterparts. No significant difference was seen in pH3+ cells between conventionally reared wild-type female flies (*w^1118^*) and axenically reared injured flies. The mitotic index of *upd2,3^Δ^* females displays reduced gut proliferation compared to wild-type (*w^1118^*) flies 8 h after wounding of axenic flies. CI: clean injury; SI: septic injury (B) Flies where *upd3* was knocked down in hemocytes after an injury, i.e., *hmlΔGAL4* > *UAS- upd3-*RNAi show an attenuation of mitotic index 8 h post-injury. (C) Nuclei staining using Lamin antibody *hmlΔGAL4* > *UAS-upd3*-RNAi was compared to the wild-type counterparts (*hmlΔGAL4* > *Cs*). Quantifying the number of nuclei per gut section of intestines stained with Lamin antibody reveals increased nuclei in fly intestines with reduced intrahemocyte ROS after an injury in *hmlΔGAL4* > *UAS-upd3-*RNAi vs the wild-type flies. (D) The mitotic index of the midgut of flies where *upd3* was knocked down in hemocytes after an injury, i.e., *hmlΔGAL4* > *UAS-upd3*-RNAi adult flies, displays increased proliferation or hyperplasia at 10 days post-injury as compared to wild-type (*hmlΔGAL4* > *w^1118^*). The increased number of mitotic stem cells was revealed by increased pH3+ immunostaining in dissected midguts from female flies. Values represent mean ± SD from three independent experiments. Statistical analyses were performed using one-way ANOVA with Tukey’s post hoc test (A, B, D) or unpaired Student’s *t*-test (C). Scale bars, 50 μm.

We next examined whether regeneration of the intestine is influenced by the activation of hemocytes following wounding at a distant site (Fig. S1A). When *upd3* expression was silenced in hemocytes using RNA interference, ISC proliferation was markedly reduced, closely resembling the phenotype of axenic *upd2,3^Δ^* mutants (Fig. 2B). Likewise, *Drpr*—required for hemocyte activation and *upd3* induction after injury (Chakrabarti & Visweswariah, 2020)—was essential for this proliferative response: *Drpr^Δ5^* mutants showed a similar decrease in mitotic ISCs (Fig. S3A). These findings establish that hemocyte-derived Upd3 is critical for triggering remote ISC proliferation after injury, with minimal contribution from microbiota-derived cues. The reduced proliferative response observed 8 hours post-injury in the absence of hemocyte-derived Upd3 correlated with later defects in epithelial integrity, as indicated by the loss of adherens junctions one-week post-injury (Fig. 1C).

Given the importance of hemocyte-derived Upd3 in early regenerative signaling, we next asked whether its absence influences long-term intestinal homeostasis, particularly under conditions associated with tissue ageing. In older flies, intestinal hyperplasia and dysregulated ISC proliferation are well-established hallmarks of age-related decline contributing to reduced lifespan (Biteau et al., 2010; Buchon et al., 2009). To access this, we examined nuclear morphology and proliferative activity following injury in flies lacking activated hemocytes. Lamin staining revealed increased nuclei in intestines from hemocyte-specific *upd3* knockdown flies and *Drpr^Δ5^* mutants seven days post-injury, relative to controls (Fig. 2C, S3B). Consistently, the mitotic index remained elevated a week after injury in flies lacking hemocyte-derived Upd3, indicating persistent hyperplasia and a failure to restore homeostasis (Fig. 2D).

In summary, our results show that in the absence of hemocyte-derived Upd3, adherens junctions are compromised and essential for maintaining epithelial integrity following injury. This junctional breakdown causes leaky enterocytes, and regenerative proliferation accelerates into hyperplasia. Together, these defects disrupt intestinal homeostasis and likely underlie the early lethality observed in flies unable to activate Upd3 expression in hemocytes after injury.

### Chronic STAT activity in the gut is critical to surviving injury by maintaining epithelial turnover

Hemocyte-derived Upd3 acts through the JAK/STAT signaling pathway to regulate epithelial renewal in the intestine. Upd3 binds to the ubiquitously expressed receptor Domeless, activating the Janus kinase Hopscotch and leading to phosphorylation and nuclear translocation of the transcription factor Stat92E (Fig. S4A). We previously demonstrated that hemocyte-derived Upd3 can activate JAK/STAT signaling in the gut following wounding, promoting intestinal regeneration and survival (Chakrabarti et al., 2016). To investigate the temporal and spatial dynamics of STAT92E activation in the gut, we used a 10XSTAT-GFP reporter, which expresses a destabilized version of GFP (Bach et al., 2007).

Reporter expression was strongly induced in the visceral muscle and enterocytes within 24 hours of thoracic injury and persisted for up to 10 days (Fig. 3A) which we quantified at day 5 post-injury, indicating sustained activation of JAK/STAT signaling during the repair phase. Activation was most prominent in the middle and posterior midgut regions (Fig. 3B), distant from the injury site, highlighting the systemic nature of this hemocyte-to-gut signaling axis.

**Fig 3.**
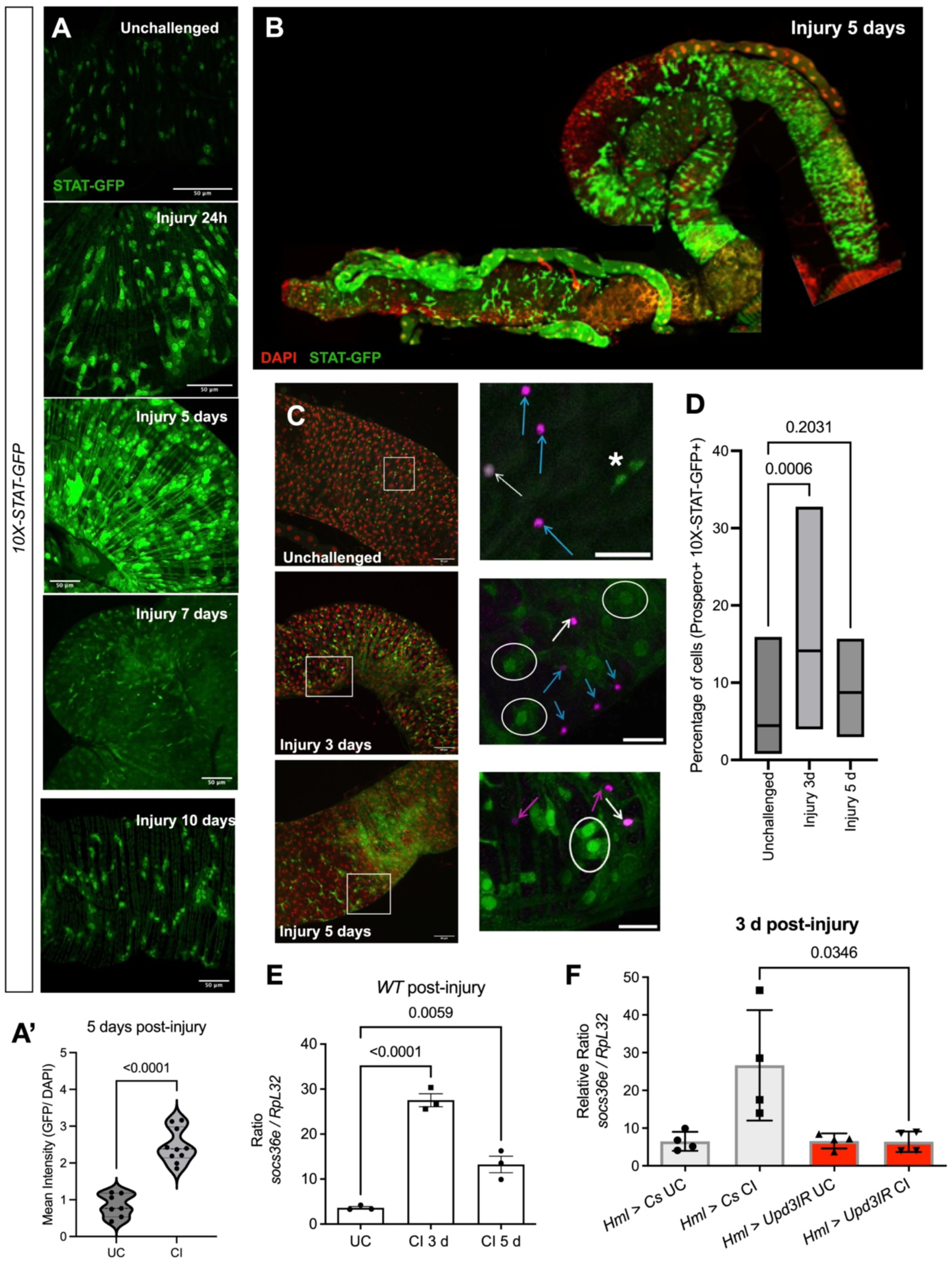
Chronic STAT activation in the intestine post-injury by hemocytes. (A) 10X-STAT-GFP reporter expression in adult female midguts showing JAK/STAT activity at different time points after sterile injury. Reporter activation appears in visceral muscles and intestinal progenitor cells by 6 hours post-injury, expands throughout the gut by day 5, and remains weakly detectable in visceral muscles at day 10. Representative images show posterior midgut regions of unchallenged (UC) and injured flies. (A’) Quantification of mean GFP intensity normalized to DAPI is shown for unchallenged and 5-day post-injury guts (UC, n = 7; 5 dpi, n = 10). (B) Whole-mount midgut of a female fly 5 days post-injury showing regionalized JAK/STAT activation, with maximal reporter activity in the middle and posterior regions. DAPI is in red and STAT92E (10X-STAT-GFP) is in green. (C) 10X-STAT-GFP reporter activity in the posterior adult midgut under unchallenged and post-injury conditions. Top panels: Unchallenged midguts (left, overview; right, magnified view) showing baseline STAT reporter activity. Middle panels: Midguts 3 days post-injury (left, overview; right, magnified) stained for Prospero, showing transient STAT activation in both enterocytes and EEs. Bottom panels: Midguts 5 days post-injury (left, overview; right, magnified) showing sustained STAT activation in enterocytes and minimal activity in Prospero-positive EEs. For all panels, asterisks (*) mark STAT-positive intestinal stem cells (ISCs). White arrows indicate Prospero-positive, STAT-GFP– positive enteroendocrine (EE) cells, blue arrows mark Prospero-positive EEs lacking STAT activity and white circles indicate STAT-GFP positive enterocytes (ECs). (D) Quantification of the total number of Prospero-positive and 10X-STAT-GFP–positive cells in the posterior midgut. A transient increase in STAT activation was observed at 3 days post-injury, which subsided by day 5, consistent with a sustained response primarily in enterocytes. (E) RT-qPCR analysis of the STAT92E target gene *socs36E* in guts from wild-type flies 72 hours post-injury shows significant induction relative to unchallenged controls. (F) In flies with hemocyte-specific knockdown of *upd3* (*hmlΔGAL4* > *UAS-upd3*-RNAi flies), *socs36E* induction in the intestine was markedly reduced compared with wild-type counterparts (*hmlΔGAL4* > *Cs*). For E-F, experiments were repeated thrice with about 20-30 flies in each experiment. P values are shown and are determined by the Student’s t-test.

To resolve the cellular pattern of STAT activation, we co-stained the midgut with Prospero to mark enteroendocrine (EE) cells and quantified 10XSTAT-GFP reporter activity at different time points after injury. STAT signaling was evident in both enterocytes and Prospero-positive EEs three days after injury, but by day five it remained strongly sustained only in enterocytes, with minimal activity in EEs (Fig. 3C). Despite this transient activation, the total number of Prospero-positive cells did not change throughout recovery (Fig. S4B). Quantitative analysis of polyploid enterocytes revealed a significant increase in Stat92E reporter activity 5 days post-injury (Fig. 3D), confirming that chronic JAK/STAT activation occurs in differentiated intestinal cells during the recovery period.

We examined the canonical JAK/STAT target gene *socs36E* expression to further validate this sustained activation. Transcript levels of *socs36E* were robustly induced 72 hours post-injury in wild-type flies (Fig. 3E), consistent with ongoing STAT activation. However, *socs36E* induction was markedly reduced in flies with hemocyte-specific *upd3* knockdown, which fail to activate hemocytes after wounding (Fig. 3F).

To investigate the role of STAT92E in epithelial turnover of the intestine post-injury, we employed the *esg^ts^ Flp-Out (esg^ts^F/O)* system. *UAS-Flp* recombinase is induced in intestinal progenitor cells by a temperature shift using *esgGal4^ts^*. As injury-induced ISC proliferation, we observed that on day 1 post-heat shock, the clones were indeed larger following injury (Fig. 4A vs 4A’, Fig. 4A’’). The size of the clones returned to normal size 6 days post-heat shock (Fig. 4B vs 4B’). As previously shown by others (Lin et al., 2010; Liu et al., 2010), the loss of STAT92E by RNAi under basal conditions led to smaller clone size and defects in enterocyte differentiation (Fig. 4B and S4D). However, knockdown of STAT92E following injury resulted in clones remaining large even 6 days post clone induction (Fig. 4B). On quantification of clone size, we found that at 6 days post heat shock in wild-type flies after injury, the clone size returns to that of unchallenged wild-type flies (Fig. 4B”). However, clones with STAT92E knocked down following injury are significantly larger in size than those seen in unchallenged flies (Fig. 4B”). Thus, STAT signaling is essential for midgut turnover in response to injury.

**Fig 4.**
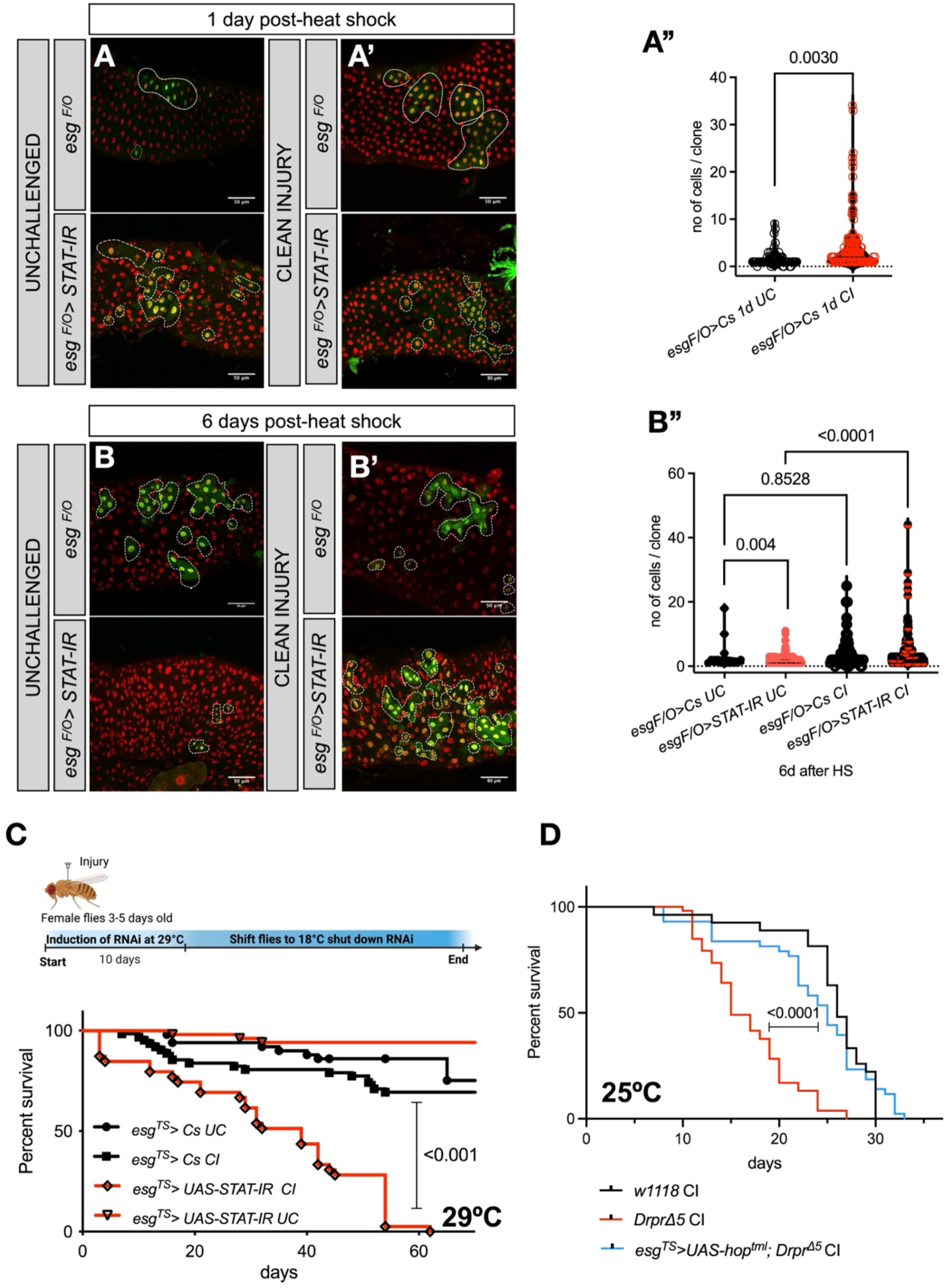
STAT92E activation in the intestine is required for the turnover of enterocytes and survival after an injury. (**A-A’”** Epithelial cell turnover was monitored at 1-day post-heat shock at (37°C for 30 mins; to activate the hs-flippase), with and without injury (A’’ and A respectively), following inhibition by *Stat92E RNAi* expressed under *Esg^ts^F/O* control (*esg-Gal4 tubGal80^ts^ UAS-GFP/CyO; UAS-Flp>CD2>Gal4/TM6B*; Jiang et al., 2009). The green cells mark the clones where the activity of driver *esgGAL4* is turned on. Nuclei are shown in red. (**A””**) There are significantly more cells in clones at day 1 post-induction in the intestines of injured animals, as determined by Student’s t-test. UC: unchallenged, and CI: clean injury. (**B-B”**) Epithelial cell turnover when *STAT92E RNAi* was expressed without injury 6 days post-heat shock, resulted in a smaller clone size comprised mostly of small non-differentiating proliferative progenitor cells. On the other hand, post-injury, when *STAT92E RNAi* was expressed without injury, resulted in EC accumulation and a diffused GFP signal in differentiated ECs. (**B”**) There are significantly fewer cells in clones at day 6 post-induction in the intestines of knockdown animals under unchallenged conditions. Clone size reaches homeostasis on day 6 post-induction, unchallenged with injured intestines. Region 2 of the intestine is shown. UC: unchallenged, and CI: clean injury. P values shown are from a one-way ANOVA with a “Tukey’s multiple comparisons test.” (C) Temporal knockdown of *STAT92E* in intestinal progenitor cells using RNAi for 10 days post-injury. Flies per genotype are pooled from at least three independent experiments. Log-rank test was used for comparing wild type *esg^TS^GAL4* > *Cs* (UC, n = 50 & CI, n = 62) and *esg^TS^GAL4* > *STAT92eIR* (UC, n = 51 & CI, n = 39). (D) Expression of constitutively active JAK in stem cells. Log-rank test used for comparing wild type (*w^1118^*; CI, n = 40) and *drpr^Δ5^* (CI, n = 53) flies as compared to *STAT92E* expression in ISCs, *esg^T^GAL4* > *UAS-Hop^tml^*; *drpr^Δ5^* n = 43). UC: unchallenged & CI: Clean Injury.

To test the relevance of chronic STAT92E activation post-injury in the gut, we reduced activation of the JAK/STAT signaling pathway by overexpressing a dominant-negative form of the receptor *Domeless* (*Dome^DN^*) in gut progenitor cells using *esg^ts^GAL4*. These knocked-down flies were more susceptible to injury (Fig. S4E). Since STAT92E was activated in the gut chronically for about 10 days (Fig. 3C-D), we blocked the pathway in the gut for the first ten days post-injury. This acute knockdown of STAT92E for 10 days increased flies’ susceptibility to injury (Fig. 4C). This means that STAT92E chronic activation post-injury is critical for flies to maintain gut homeostasis and their survival.

To test whether chronic activation of STAT92E in the gut could compensate for the reduced induction of Upd3 in hemocytes, we activated the JAK/STAT pathway specifically in the intestines of flies that fail to induce Upd3 and cannot activate STAT92E in any tissue upon wounding. To this end, we overexpressed a constitutively activated version of JAK called *hop^Tum-l^* in the gut progenitor cells using *esg^ts^GAL4* in the *Drpr^Δ5^*mutant animals. *Drpr^Δ5^* mutant animals were rescued when STAT92E was activated in the gut post-injury (Fig. 4D). Similarly, only activating STAT92E in enterocytes using the enterocyte driver *NP1Gal4* rescued *Drpr^Δ5^* mutant animals from injury-induced lethality (Fig. S4F). Hence, STAT92E activation in the gut is critical to surviving an injury.

### Hemocytes home to the gut post a distant injury and protect from subsequent chronic oral infections

In mammals, the gut is the largest immune organ in the body (Mowat & Agace, 2014). However, how circulating hemocytes communicate with the intestinal epithelium in *Drosophila* remains poorly understood. We hypothesized that a subset of *upd3*-producing hemocytes might migrate to the gut after injury to sustain STAT92E activation in enterocytes.

To test this, we imaged intact intestines immediately after dissection, minimizing tissue disruption by rapid fixation. Following thoracic injury, a distinct population of hemocytes was in close contact with the middle midgut (R3–R4 region), and these cells persisted at the exact location up to 5 days post-injury (Fig. 5A–B; Fig. S5A). Quantification of gut-associated hemocytes revealed a significant increase in the number of hemocytes adhering to the midgut 3 days after injury compared with uninjured controls (Fig. 5C).

**Fig 5.**
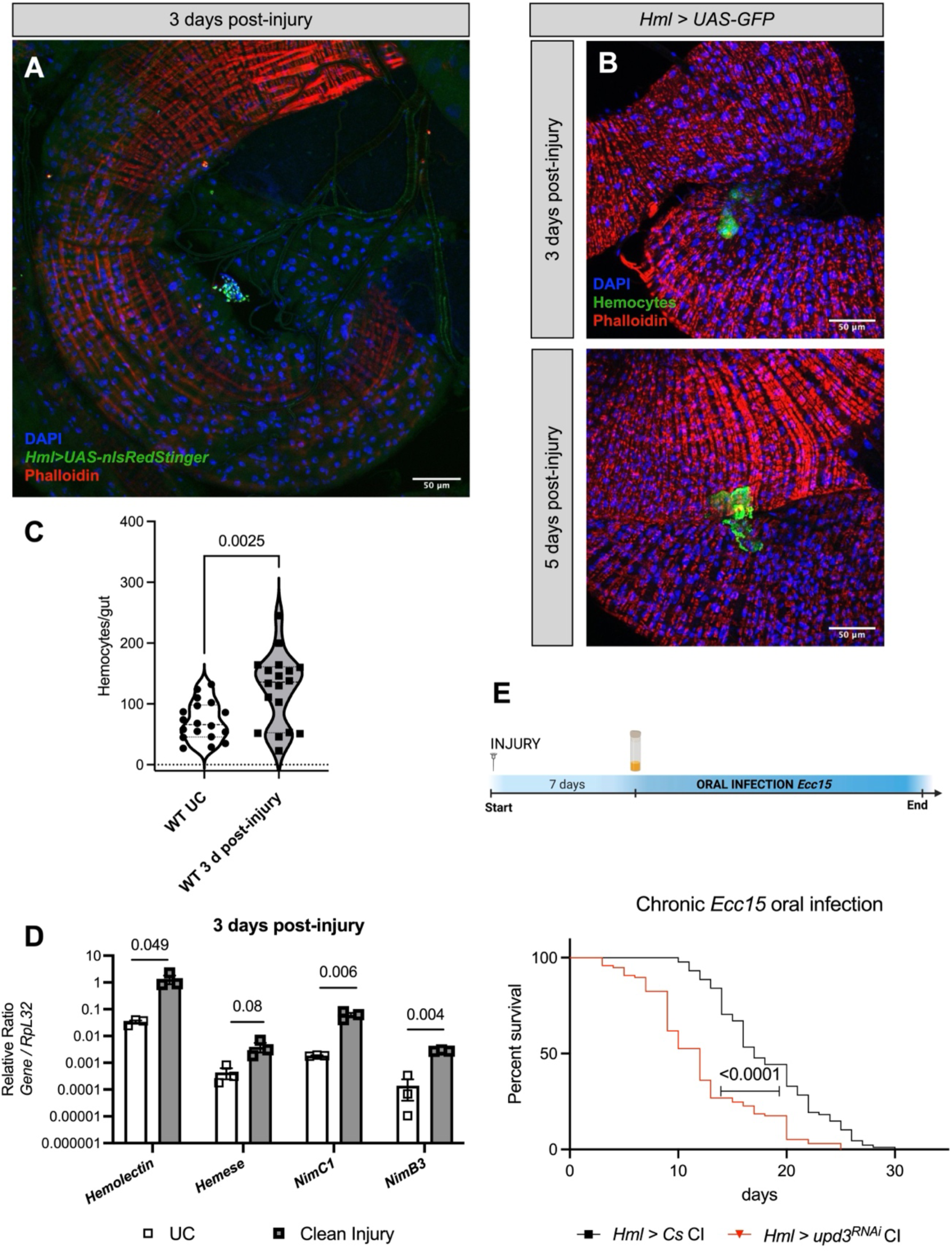
Hemocytes home to the intestine post-injury and are crucial to survive infection. (A) *hmlΔGAL4 >UAS-nlsRedStinger* fly intestines were dissected 3 days post-injury. The homing of hemocytes is seen predominantly in the R3-R4 loop of the middle midgut. (**B**) Kinetics of localization of hemocytes to the middle-midgut in adult Drosophila flies. (B) *hmlΔGAL4 >UAS-GFP* fly intestines were dissected 3- and 5-days post-injury. A representative image of 10 dissected and stained guts is shown for (A & B). (C) Hemocytes that recruit to the gut are quantified from unstretched intestines of WT (*hmlΔGAL4* > *UAS-GFP*) female flies, 72 hours with or without injury. Quantifying hemocytes attached to the entire midgut at 72 hours post-injury reveals a significant increase in recruitment. The p-value is from the Student’s t-test. (D) The hemocyte homing to the gut was assessed by employing four hemocyte marker genes: *Hml*, *Hemese*, *NimC1*, and *NimB3,* in dissected intestine post-injury. All genes were upregulated in the fly intestines 72 hours post-injury. For (D), values represent mean ± SD from experiments performed in biological triplicate. For (D), the P values shown are from a one-way ANOVA with a “Tukey’s multiple comparisons test” (**E**). Survival of flies following oral infection with *Erwinia caratovora caratovora 15* (*Ecc15*), seven days post-injury, as indicated in the scheme above, wild-type and *hmlΔGAL4* > *UAS-Upd3-*RNAi animals. The survival analysis shows that the absence of Upd3 from hemocytes post-injury makes animals more susceptible to a benign oral infection with *Ecc15*. Flies per condition are pooled from at least three independent experiments. Log-rank test used for comparing wild type (*hmlΔGAL4* > *Cs*, n = 88 & *hmlΔGAL4* > *UAS-upd3-*RNAi, n = 97).

To further confirm that hemocytes get recruited to the gut after distant injury, we measured the expression of four hemocyte-specific genes—*hemolectin (Hml), hemese, NimC1,* and *NimB3*—in dissected intestines 72 h post-injury. All four hemocyte markers were significantly upregulated relative to uninjured flies, indicating robust hemocyte association with the intestine following wounding (Fig. 5D).

We next examined whether this recruitment required hemocyte-derived Upd3. In intestines from *hmlΔ-GAL4 > UAS-upd3-RNAi* flies, hemocytes were markedly reduced or absent 3 days post-injury compared with controls (Fig. S5B). P1 immunostaining, which labels mature hemocytes, confirmed this loss: *hmlΔ-GAL4 > UAS-upd3-RNAi* flies displayed significantly fewer P1⁺ hemocytes associated with the gut 72 h post-injury than wild-type flies (Fig. S5C). Consistently, RT-qPCR analysis of *NimC1* transcripts in intestinal tissue revealed reduced expression in *hmlΔ-GAL4 > UAS-upd3-RNAi* animals after injury, confirming defective hemocyte recruitment (Fig. S5D).

Previous studies have shown that hemocytes are recruited to the gut following direct intestinal damage (Ayyaz et al., 2015). We wondered whether a distant wound that does not directly damage the gut but still leads to hemocyte recruitment could influence gut function and affect susceptibility to subsequent challenges. We therefore asked whether hemocyte recruitment after a distant wound— without direct gut injury—could influence gut resilience to future infections. To test this, we orally infected wild-type and *hmlΔ-GAL4 > UAS-upd3-RNAi* flies with *Erwinia carotovora carotovora 15* (*Ecc15*) (Basset et al., 2000) seven days after clean injury. Flies lacking hemocyte-derived Upd3 were significantly more susceptible to this otherwise non-lethal oral infection (Fig. 5E). Similarly, *Drpr^Δ5^*mutants, which fail to induce Upd3 after wounding, exhibited increased susceptibility to *Ecc15* infection when challenged seven days after injury (Fig. S5E).

Because the loss of hemocyte recruitment is accompanied by disrupted adherens junctions and compromised enterocyte integrity (Fig. 1E), we propose that the failure of hemocytes to home to the intestine weakens epithelial barrier maintenance, thereby sensitizing injured flies to secondary enteric infections. When this cross-talk between tissues is disrupted, the intestinal epithelium fails to renew, the barrier weakens, and flies become prone to secondary infections. Hemocytes thus serve a dual role— defending against damage while orchestrating the physiological link between injury response, gut integrity, and host survival (Fig 6).

**Fig 6.**
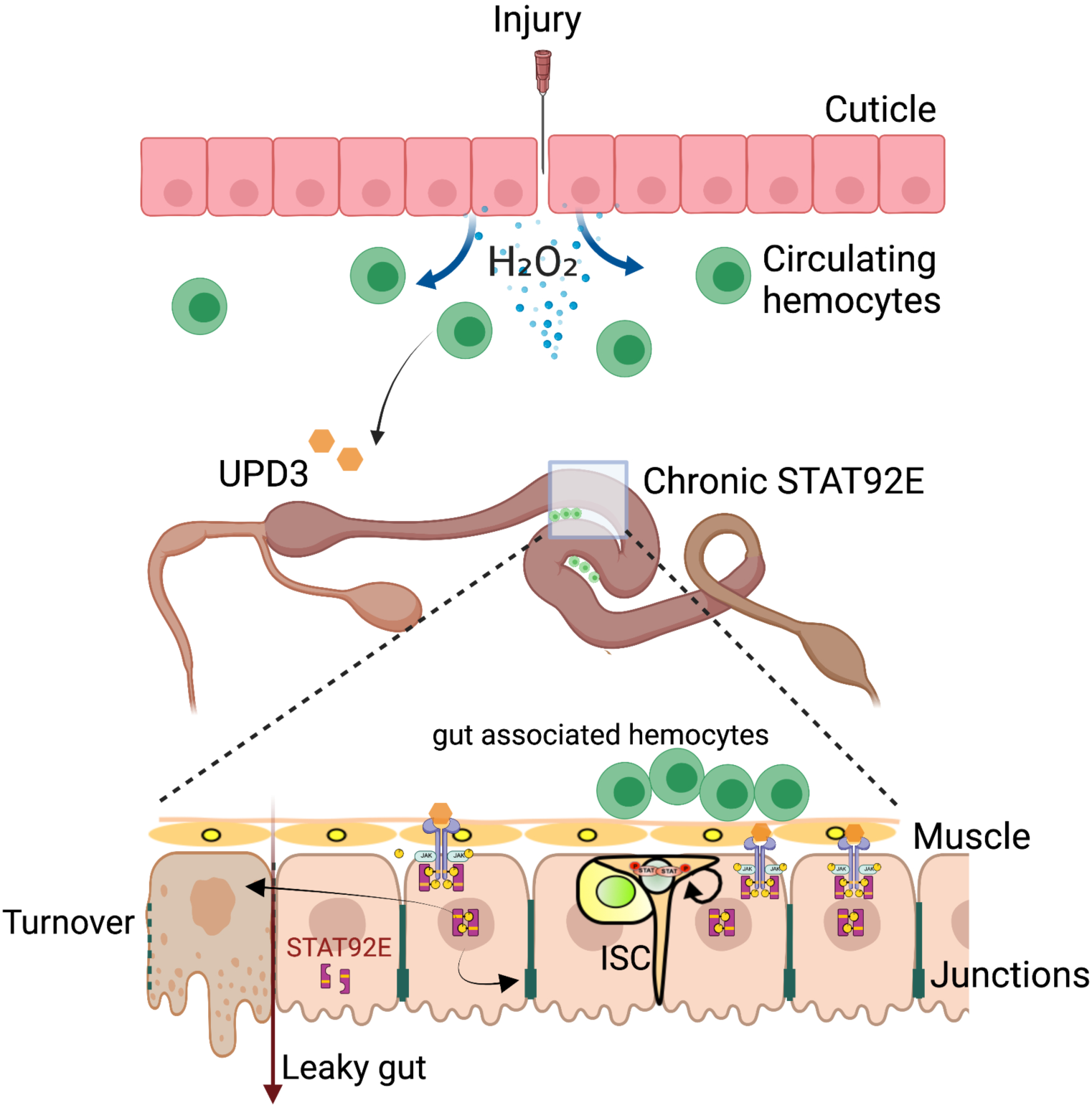
Schematic on the role of hemocytes in maintaining gut barrier function post-injury. After an injury, hemocytes sense hydrogen peroxide and release Upd3, which activates the STAT92E systemically in other organs. In the intestine, STAT92E remains chronically activated for up to 10 days following the injury and plays a vital role in maintaining gut barrier functions and gut turnover. In the absence of STAT92E activation, the junctions between enterocytes are compromised, resulting in increased intestinal permeability. Hemocytes that migrate to the gut after injury may contribute to the sustained activation of STAT92E.

## Discussion

We show here that during the wounding response, communication between the site of injury and the gut is critical for host survival. This study builds on our previous work showing that hydrogen peroxide produced at the wound activates JAK/STAT signaling in hemocytes, leading to Upd3 secretion, which is essential for survival after injury. (Chakrabarti et al., 2016; Chakrabarti & Visweswariah, 2020). This establishes a systemic axis through which signals originating at a local injury site are transmitted to distal organs such as the gut.

The regulation of gut integrity is central to organismal health, as the intestine serves as the interface between an animal and its environment (Zilbauer et al., 2023). Intestinal epithelial cells perform various critical functions in the gut, such as absorbing nutrients, maintaining a healthy microbiota balance, training the immune system, and eliminating pathogens Here, we identify a role for hemocytes in triggering intestinal stem cell (ISC) proliferation following distant injury, thereby maintaining epithelial turnover and barrier function. Comparable roles for macrophages in promoting epithelial repair have been documented in salamanders, zebrafish, and mammals (Arinda et al., 2022). Other studies have shown that wounding can remotely control caspase activity in enterocytes, which activates the intestinal stem cell regeneration pathway in the gut to dampen the otherwise lethal systemic wound reaction (Ayyaz et al., 2015; Takeishi et al., 2013). Our data extend this paradigm by demonstrating that hemocyte-derived Upd3 acts as a key mediator of this remote epithelial response.

While the molecular mechanisms maintaining gut integrity with age remain incompletely understood, our findings identify JAK/STAT signaling as a central component of the systemic wound response. Hydrogen peroxide activates hemocytes to secrete Upd3, stimulating JAK/STAT signaling in the gut. Without hemocyte-derived Upd3, flies show defects in adherens junctions, impaired barrier function, and early lethality (Figs 1 and 2). The gut barrier defects in these animals mirror those seen in age-onset intestinal permeability and inflammatory diseases such as IBD and celiac disease (Marchiando et al., 2010). We have shown that flies without hemocytes or hemocytes that do not secrete Upd3 are susceptible to a simple injury. The cause of early death in these flies could be a gut barrier dysfunction, as the same phenotype is linked with age-onset death in flies (Rera, Clark, et al., 2013). How hemocytes influence a robust localization of adherens junction proteins remains unknown. We propose that STAT activation in enterocytes and visceral muscle promotes junctional stability, as wild-type flies show a post-injury increase in junctional proteins such as Dlg1 and E-cadherin (Fig. 1). How exactly JAK/STAT signaling reinforces junctional organization remains to be explored. Notably in the Drosophila blood–brain barrier (BBB), similar mechanisms operate where JAK/STAT signaling regulates barrier integrity and macrophage invasion (Kim et al., 2021) (Winkler et al., 2025). Thus, our work and these studies highlight a conserved role for JAK/STAT signaling in coordinating immune input with epithelial barrier maintenance across tissues.

Previous work has shown that mild and severe intestinal damage elicit different regenerative outcomes—mild damage promotes repair without apoptosis. In contrast, severe injury leads to epithelial cell death and renewal through apoptosis (Loudhaief et al., 2017). Chronic activation of STAT in the gut promotes epithelial turnover, likely through apoptosis and replacement of damaged cells. The absence of STAT92E in the progenitor cells results in defective differentiation and accumulation of older enterocytes, consistent with impaired barrier function and hyperplasia (Figs 3 and 4). Thus, STAT signaling seems to be implicated in balancing proliferation, differentiation, and turnover following an injury, making sure that intestinal regeneration is proportional to the degree of systemic stress.

The presence of myeloid cells in the gut could have been the evolutionary basis for developing new organs, such as the spleen. No specific organs exist in adult insects where blood cells can monitor host antigens. The observation of hemocytes homing and attaching to the gut could resemble a primitive spleen found in lower vertebrates like hagfish, where myeloid cells are sparsely distributed along the intestine (Brendolan et al., 2007; Tischendorf, 1985). This may provide an evolutionary advantage that led to the development of organs like the spleen. Our data support this notion, demonstrating that hemocytes migrate to the gut in response to distant injuries and provide protection against ongoing oral infections. In the future, exploring the mechanisms by which activated hemocytes migrate and establish residency in the intestine for up to a week would be interesting. The central part of the midgut, characterized by a stereotypical looping in its 3D organization, exhibits the highest number of hemocytes attached to the gut. While this region is typically quiescent regarding ISC proliferation (Marianes & Spradling, 2013), injury prompts these ISCs to undergo mitosis. How this region is critical for the proper functioning of the entire gut needs to be investigated. In addition, further research is necessary to understand how hemocytes migrate and their role in gut residency and protection.

## Materials and Methods

### *Drosophila* stocks and rearing under conventional and axenic conditions

*Canton^S^* (*Can^S^*) and *w^1118^* flies were used as wild-type controls. The following fly lines were used in this study: *w; UAS-upd3-IR* (Agaisse et al., 2003), *hml*Δ*Gal4, UAS-GFP* (Sinenko & Mathey-Prevot, 2004), *Drpr^Δ5^* (Freeman et al., 2003) backcrossed, and Ubi DE-cadherin-GFP (E-Cad::GFP; from M. Narasimha, TIFR Mumbai). All RNAi strains have been validated previously in our last two studies (Chakrabarti et al., 2016; Chakrabarti & Visweswariah, 2020).

For RNAi (IR) studies, F1 progeny carrying one copy of the driver as well as one copy of the *UAS-IR* were raised at 18°C during their larval and pupal development and then moved to 29°C for 8 days to activate the *UAS-IR* 3 days post-eclosion. *Drosophila* stocks were maintained using standard fly medium comprising 8% cornmeal, 4% sucrose, 2% dextrose, 1.5% yeast extract, 0.8% agar, supplemented with 0.4% propionic acid, 0.06% orthophosphoric acid, and 0.07% benzoic acid. Unless otherwise stated, all stocks were maintained at 25°C on a 12 h light/ 12 h dark cycle.

Axenic flies were generated by bleaching embryos using 0.6% sodium hypochlorite. Embryos were washed in bleach for 3 minutes twice, followed by a single 70% ethanol wash and the final 3 washes with sterile MilliQ water. Finally, the embryos were transferred with the help of a sterile brush on autoclaved food. Sterile fly food was made by autoclaving at 121 °C and 15 psi for 30 min and then immediately put on a horizontal shaker to prevent separation during cooling. Filter-sterile preservatives were added under sterile conditions before the food was cooled. The presence of bacteria in gut homogenates was examined by PCR amplification of 16S rRNA genes using eubacterial primers (27F and 1492R) and by culturing the homogenates on mannitol agar or 1/10-strength tryptic soy agar.

### Injury and infection experiments

For a clean injury, flies were pricked in the thorax under the wing with a tungsten needle. For injury of axenic flies, the needle was sterilized by flaming, followed by UV treatment for 15 minutes. All injections of axenic flies were done under a laminar hood using ice to anesthetize flies.

*Erwinia carotovora carotovora 15 (Ecc15)* is a Gram-negative bacterium described by (Basset et al., 2000). *Ecc15* was cultured overnight in Luria broth at 29°C, and *E. faecalis* was cultured overnight at 25°C. For septic injury experiments, *Drosophila* 3-4 days old adults were used. For oral infection, flies were starved for 2 hours and then fed a mixture of concentrated culture of *Ecc15* (OD600 ∼200) with 5% sucrose and shifted to 29°C for optimal bacterial proliferation. Flies in survival experiments were kept on a medium without fresh yeast, and survivors were counted daily.

### Smurf Assay

We referred to (Rera et al., 2012) for the Smurf assay conditions. The blue dye medium was prepared using a standard medium with Brilliant blue FCF (Sigma-Aldrich, final 2.5% wt/vol) to prepare the feeding medium. Flies were fed with this medium at 25°C for 1 day, then transferred to a new vial containing cornmeal-agar food without blue dye to clean the epidermis before classifying the Smurf level.

### Survival assay

Flies of the appropriate genotypes were collected within 1–2 days after hatching at 18°C and kept in groups of 20 flies per vial. The flies were transferred to 29°C with a 12 h light/ 12 h dark cycle. The number of dead flies was recorded every other day. The media was changed every 3-4 days. The Kaplan-Meier log-rank test was used to assess statistical significance, and the data were plotted using GraphPad Prism 9.5.

### Propidium iodide feeding

To assess intestinal barrier integrity using a small-molecule tracer, we employed propidium iodide (PI; molecular weight ∼668 Da). Under normal conditions, PI does not cross intact epithelial junctions when administered orally. In guts with compromised barrier function, PI enters enterocytes and accumulates in the cytoplasm, providing a sensitive readout of epithelial leakiness.

For the assay, adult flies were starved for 2 h and then fed a solution of 100 μM PI in 5% sucrose for 2 h. Guts were dissected immediately, mounted in VECTASHIELD Antifade Mounting Medium with DAPI (Vector Laboratories, H-1200-10), and imaged using an Olympus FV3000 confocal laser scanning microscope with a 40× NA 1.4 oil objective.

### RT-qPCR

Around 30 fly intestines were dissected and collected in 500μl of Trizol (Takara Bio, 9108). Total RNA was extracted according to the manufacturer’s instructions. The quality and quantity of RNA were determined using a NanoDrop ND-1000 spectrophotometer (Thermo Fisher). 1μg of RNA was treated with 1U of DNase I (Thermo Fisher, EN0521), following which cDNA was generated using Revertaid reverse transcriptase (Thermo Fisher, EP0441). RT-qPCR was performed using dsDNA dye SYBR Green (Takara Bio, RR420A) on a Biorad CFX96 real-time PCR machine. Expression values were normalized to *RpL32*.

Primers used in this study were as follows: *RpL32* forward 5′-GACGCTTCAAGGGACAGTATCTG- 3′, *RpL32* reverse 5′-AAACGCG-GTTCTGCATGAG-3′; *AttD* forward, 5’’ AAGGGAGTTTATGGAGCGGTC -3’’ *AttD* reverse, 5’-GCTCTGGAAGAGATTGGCTTG -3’’ *upd3* forward 5’-GCGGGGAGGATGTACC, *upd3* reverse, 5’-GTCTTCATGGAATGAG-CC-3’’ *socs36E* forward, 5’-GCACAGAAGGCAGACC-3’’, *socs36E* reverse, 5’-ACGTAGGAGACCCGTAT -3’’

### Imaging and immunohistochemistry

For immunofluorescence, 3 to 5-day-old females were dissected in 1X PBS, fixed for 20 minutes in PBS 0.1% Tween 20 (PBT), and 4% paraformaldehyde. Samples were incubated in a blocking solution (1×PBS containing 2% BSA and 0.1% Triton X-100) for one hour and then stained with the appropriate primary antibody. The following primary antibodies were used: mouse anti-Discs large (Developmental Studies Hybridoma Bank (DSHB) #4F3, 1:100), rabbit anti-phospho-Histone H3 (Ser10) (Thermo Fisher #701258, 1:1000), and mouse anti-Lamin (DSHB #ADL67, 1:100) in PBT + 2% BSA and incubated overnight at 4°C. Subsequent washing (3 times for 20 mins) and incubation with secondary antibody were done in PBT. Secondary staining was performed with Alexa 488/594 antibodies 1:1000 (Molecular Probes/Invitrogen). Both primary and secondary antibodies were diluted in the blocking solution. DNA was stained with a 1/15000 dilution of 4’’6-diamidino-2-phenylindole DAPI (Sigma, D9542). The stained gut tissue was mounted in VECTASHIELD Antifade Mounting Medium (Vector Laboratories). PH3-positive cells were counted along the gut with an Axioplot imager (Zeiss). For imaging, gut sections were imaged using an Olympus FV 3000 confocal laser scanning microscope (CLSM) using a 40× NA 1.4 oil objective. Image panels were compiled using Photoshop (Adobe).

To visualize gut-associated hemocytes, intestines were dissected and fixed in situ to preserve adherent P1⁺ cells. Briefly, 3–5-day-old adult flies were dissected directly into 4% paraformaldehyde in PBS, leaving the dorsal abdomen intact, and fixed for 1 h at room temperature without agitation. Following fixation, non-intestinal tissues were gently removed, and the gut was dissected with its natural looping preserved. Guts were permeabilized with 0.1% PBST (PBS containing 0.1% Tween 20) for three 10-minute washes, blocked in 2% BSA for 45 minutes, and incubated overnight at 4°C with mouse anti-P1 antibody (DSHB, 1:30 in 2% BSA). After washing, samples were incubated with Alexa Fluor 555–conjugated anti-mouse secondary antibody (1:1000, Molecular Probes) for 2 h in the dark, followed by nuclear staining with DAPI (1:15,000 in PBST). Guts were mounted in ProLong™ Diamond Antifade Mountant with DAPI (Thermo Fisher Scientific) and imaged on a Nikon confocal microscope using identical settings for all genotypes.

### Quantification of PH3-positive ISCs and STAT-GFP quantification

For PH3 staining, 10-15 digestive tracts were stained with rabbit anti-PH3 antibodies, as described above. The whole midgut (from the bottom of the proventriculus to the top of the pylorus, the hindgut-midgut transition area) was scanned using an Axio-Observer Z1 microscope (Carl Zeiss Microimaging) to count the number of PH3-positive cells. The experiment was performed independently at least 3 times to statistically compare the average number of PH3-positive cells per gut.

GFP and DAPI channels from the gut region are split in Fiji. ROIs containing GFP-positive and corresponding DAPI+ cells are selected using the draw tool and added to the ROI Manager for each Z-stack. Mean GFP intensity is normalized to DAPI by calculating the GFP/DAPI ratio. The average of all Z-stack ratios for each gut is taken as the final mean intensity value.

For quantification of Prospero-positive enteroendocrine (EE) cells that were STAT active, intestines from unchallenged, 3-day post-injury, and 5-day post-injury flies carrying the 10XSTAT-GFP reporter were dissected and immunostained with mouse anti-Prospero antibody (DSHB, 1:100) as described above. Prospero-positive EE cells were identified based on their small, diploid nuclei and characteristic morphology, which distinguish them from polyploid enterocytes. The total number of Prospero⁺ cells and the number of Prospero⁺/10XSTAT-GFP⁺ double-positive cells were quantified for each gut using Fiji. The percentage of STAT-active EEs (Prospero⁺/STAT⁺) was calculated for each gut, and mean values from 10–15 intestines per condition were plotted using GraphPad Prism.

### Quantification of Dlg enrichment at bicellular junctions

To measure and compare the Dlg enrichment at bicellular junctions relative to cytoplasmic levels, we selected the bicellular junction between two cells using ROI function in Fiji. Analyze → Tools → ROI manager → Select (area of interest) → Save. After selection of ROI, we measured the fluorescence intensity of Dlg at bicellular junction. Using the same ROI dimensions, the ROI was shifted to measure the cytoplasmic fluorescence of Dlg in the cells forming the junction. The fluorescence intensity ratio was calculated by dividing the bicellular junction intensity with cytoplasmic intensity to get the relative ratio of Dlg enrichment. This was repeated for 20 cells in the R4 region of the gut and done for 3 to 4 gut samples.

### Statistics

Each experiment was repeated independently a minimum of three times (unless otherwise indicated), and error bars represent the standard deviation of replicate experiments (unless stated otherwise). All data were assumed to be parametric, and statistical significance was determined using Student’s t-test, one-way ANOVA, or log-rank test on GraphPad Prism 9.5, and P values of <0.05, <0.01, and <0.001 were considered significant. All P values are represented in each figure.

## Acknowledgments

We thank T. Kuraishi, Bloomington Drosophila Stock Center, National Institute of Genetics, Vienna VDRC, and the National Institute of Genetics for fly stocks. We thank Namyashree Nayak, Shreya H, and Janani for technical help and Soumyadip Sarkar for reading the manuscript. We are grateful to Irene Miguel-Aliaga for fruitful discussions and guidance.

This work is supported by the grant from the Wellcome Trust-DBT India Alliance (grant no. IA/I/23/2/506985 and IA/E/15/1/502342) to S. Chakrabarti. S. Visweswariah is a JC Bose National Fellow (SB/S2/JCB-18/2013), a National Science Chair supported by SERB ((NSC/2022/000019) and a Margdarshi Fellow supported by the Wellcome Trust DBT India Alliance (IA/M/16/1/502606). Y.S. Sheregare is supported by the Manipal Academy of Higher Education for the Dr. TMA Pai fellowship.

## Author contributions

S.C. conceived the project. S.C. and Y.S.S. performed the experiments. S.C. and Y.S.S. formal analysis. S.C. and Y.S.S. investigation. S.C. wrote the original draft. S.C. and S.S.V. discussed the results. S.C., Y.S.S., and S.S.V. revised the manuscript.

