## Supplemental Figures for "Drosophila blood cells bridge distant injury and gut homeostasis through Upd3-mediated inter-organ signaling"

### Supporting information

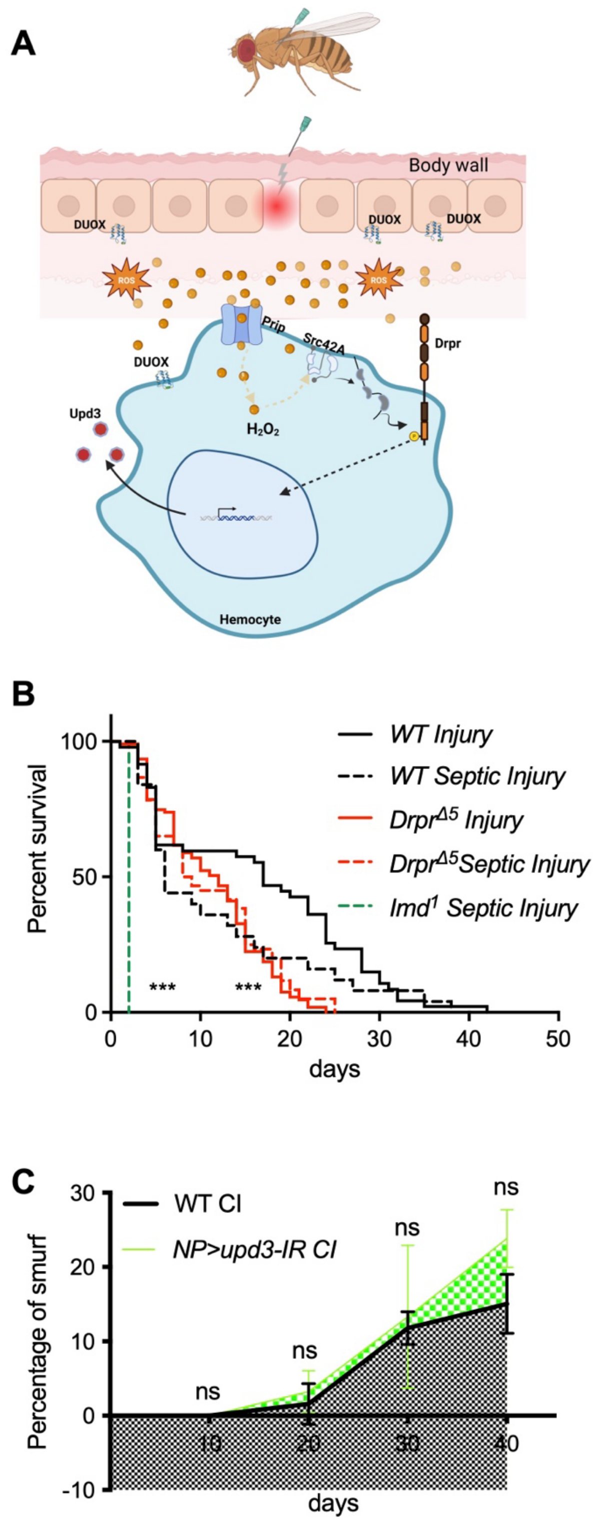


**S1 Fig. Hemocytes are important to resist injury.**

(**A**) Schematic to show signaling events in hemocytes following an injury. Hemocytes respond to the damage signal $H_2O_2$ through the kinase Src42A and its downstream target, Draper. The aquaporin channel Prip increases intracellular ROS levels to trigger the production of UPD3 cytokine from hemocytes.

(**B**) The lifespan of *Draper* knockout ($Drpr^{\Delta 5}$) flies, and *immune deficiency* ($imd^{1}$) flies susceptibility to injury and septic injury compared to the wild type ($w^{1118}$). Flies per genotype are pooled from at least three independent experiments. Log-rank test used for comparing wild type ($w^{1118}$; CI, n = 47 & SI, n = 25) and $drpr^{\Delta 5}$ (CI, n = 107 & SI, n = 60) flies and $imd^{1}$ (SI, n = 28). UC: unchallenged; CI: Clean Injury; SI: Septic Injury. Log-rank test was used to compare wild type ($w^{1118}$)

(**C**) Smurf assay with blue dye. The curve represents the cumulative proportion of dead Smurfs in the population. Flies with reduced *upd3* expression in enterocytes using the *NP1-GAL4* (*NP1-GAL4*) driver line (green curve), i.e., *NP1> UAS-upd3IR*, display same proportion of Smurf phenotype after clean injury (CI) as compared to their wild-type counterparts (grey curve).

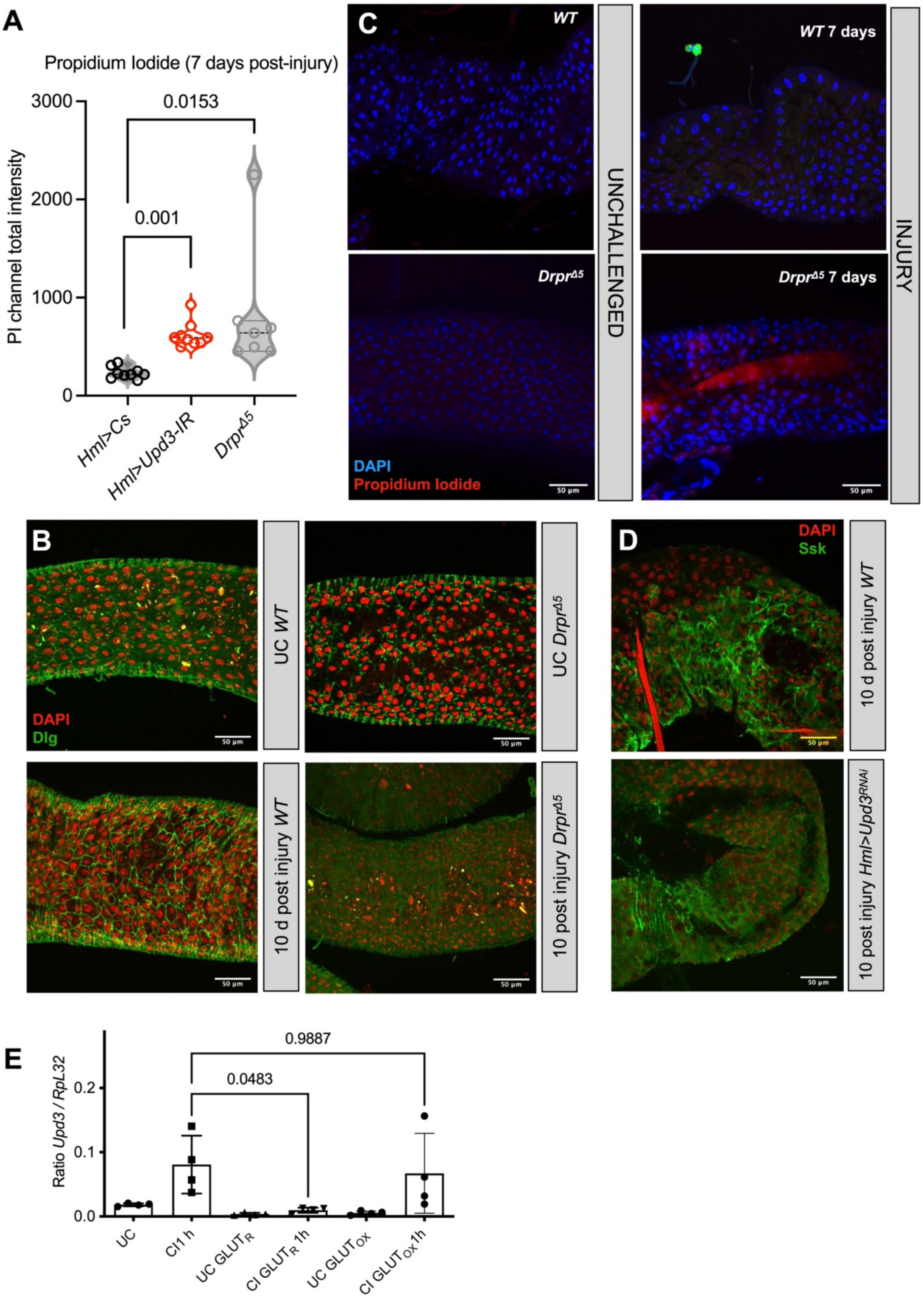


**S2 Fig. Hemocytes are essential for maintaining the intestinal barrier.**

A) Quantification of the propidium iodide (PI) feeding assay in control (*hmlΔ-GAL4* > C*S*) and hemocyte-specific *upd3* knockdown (*hmlΔ-GAL4* > *UAS-upd3*-RNAi) flies and *Drpr*$^{\Delta5}$ mutants.

Knockdown animals and *Drpr*$^{\Delta 5}$ mutants displayed significantly increased PI uptake, consistent with barrier leakiness.

(B) Midgut epithelium stained for DAPI (red) and Discs large (Dlg, green). Compared to wild type (*w*$^{1118}$), *Drpr*$^{\Delta 5}$ mutants exhibited loss of epithelial junctions 10 days post-injury.

(C) PI feeding assay revealed increased epithelial permeability in *Drpr*$^{\Delta 5}$ mutants compared to wild type (*w*$^{1118}$).

(D) Midguts stained for Snakeskin (Ssk, green) and DAPI (red) showed loss of septate junction staining, and more cytoplasmic staining in *hmlΔ-GAL4* > *UAS-upd3*-RNAi animals 10 days after injury, relative to control (*hmlΔ-GAL4* > *CS*).

(E) qRT-PCR analysis of *upd3* expression in hemocytes isolated 1 h after injury. Reduced glutathione feeding significantly suppressed *upd3* transcript levels, whereas oxidized glutathione feeding did not affect *upd3* expression. Data confirm that reduced glutathione feeding diminishes hemocyte *upd3* induction post-injury.

Statistical tests: Student's *t*-test (A, C, E) or one-way ANOVA with post hoc test where indicated. Values represent mean ± SD from at least three independent experiments. Scale bars, 50 µm.

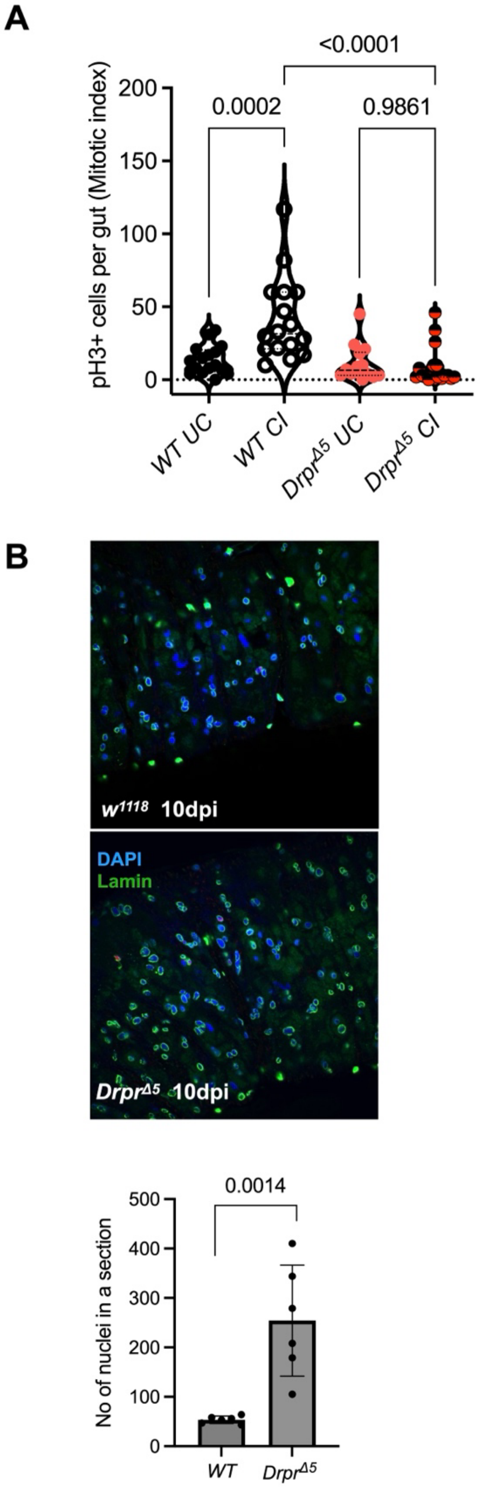


**S3 Fig. Loss of production of Upd3 from hemocytes impacts the intestinal homeostasis**.

(**A**) The mitotic index (as measured by Phospho-Histone-3- positive cells (pH3+) of the midgut flies with reduced intrahemocyte ROS after an injury in *Drpr*$^{\Delta 5}$ mutants and WT flies 8h post-injury.

(**B**) Quantifying the number of nuclei per gut section of intestines stained with Lamin antibody reveals increased nuclei in fly intestines with reduced intrahemocyte ROS after an injury in $Drpr^{\Delta 5}$ mutants. Values represent mean ± SD from experiments performed in biological triplicates. P values shown are from a one-way ANOVA with a "Tukey's multiple comparisons test."

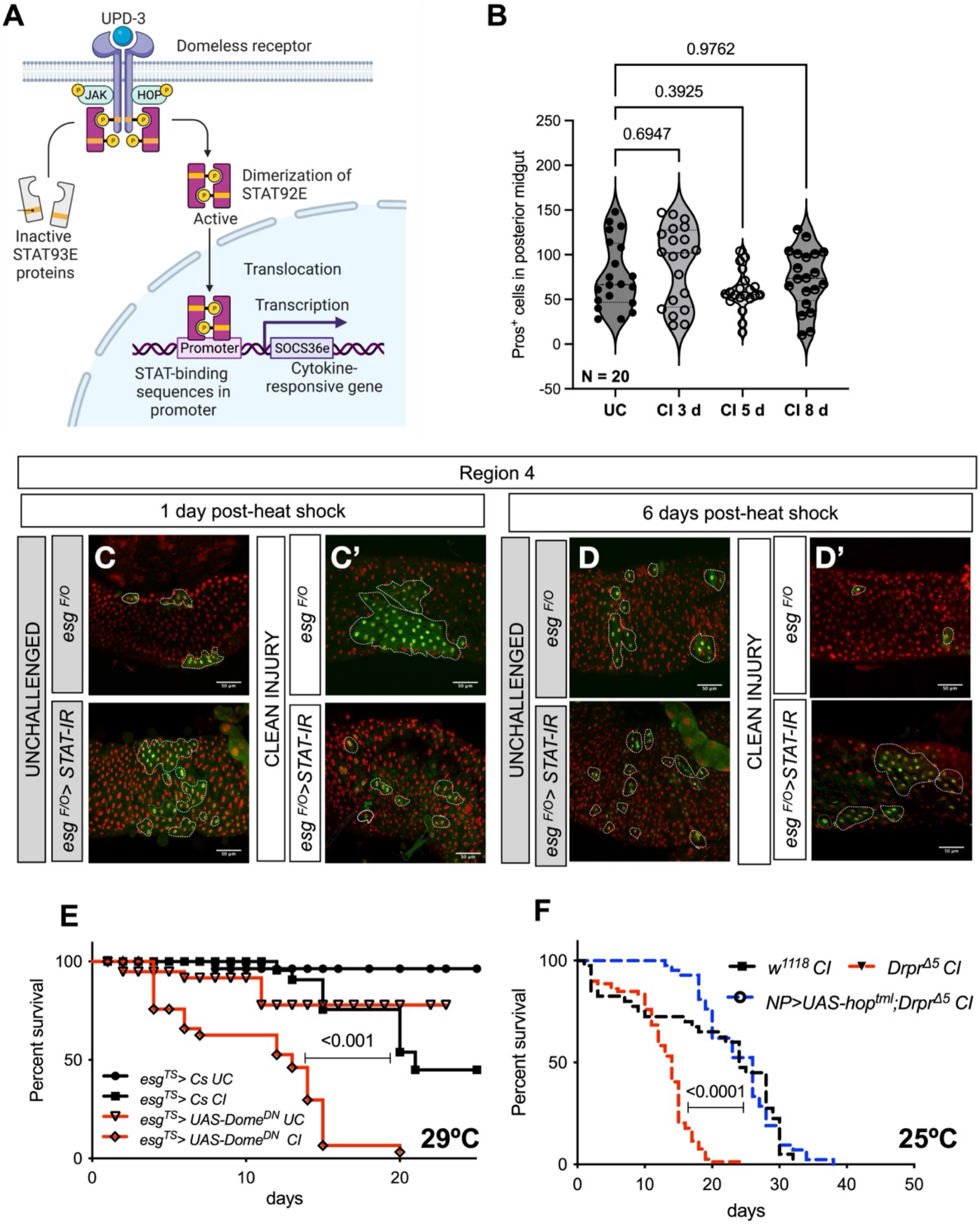


**S4 Fig. STAT92E activation in the gut requires hemocytes.**

(A) Schematic of JAK/STAT pathway activation post-injury. Hemocyte-derived Upd3 binds the receptor Domeless, leading to phosphorylation of the transcription factor Stat92E by the Janus kinase Hopscotch. Activated Stat92E translocates to the nucleus and induces transcription of its target genes.

(B) Quantification of enteroendocrine (EE) cells using Prospero staining. No significant difference in Pros+ cell numbers was detected between uninjured controls and guts examined 3, 5, or 8 days post-injury ($n$ = 20 per condition).

(C–C′) Lineage tracing using the *esg*$^{ts}$*F/O* system (*esg-Gal4 tubGal80*$^{ts}$ *UAS-GFP/CyO; UAS-Flp>CD2>Gal4/TM6B*; Jiang et al., 2009) to knock down *Stat92E* in progenitor cells. At 1 day post–heat shock (37 °C, 10 min), uninjured guts generated large GFP+ clones, whereas *Stat92E*-RNAi clones were reduced in size and composed mainly of small, undifferentiated progenitors.

(D–D′) When lineage tracing was combined with clean injury, *Stat92E* knockdown clones displayed accumulation of enterocytes (ECs) with diffused GFP signal, in contrast to control clones. This indicates that Stat92E is required for proper differentiation of progenitors after injury.

(E) Overexpression of dominant-negative Domeless (DomeDN) in progenitors using the esg^ts driver significantly increased mortality after clean injury. Survival curves: *esg*$^{TS}$*GAL4 > CS*, UC ($n$ = 41), CI ($n$ = 44); *esg*$^{TS}$*GAL4 > UAS-Dome*$^{DN}$, UC ($n$ = 40), CI ($n$ = 36). Data were pooled from three independent experiments and analyzed by log-rank test.

(F) Enterocyte-specific expression of constitutively active JAK (*Hop*$^{tum-l}$) rescued the lethality of *drprΔ5* mutants after injury. Survival curves: wild type (*w*$^{1118}$), CI ($n$ = 40); ; *Drpr*$^{Δ5}$, CI ($n$ = 79); *NP1GAL4 > UAS-Hop*$^{tml}$; *Drpr*$^{Δ5}$, CI ($n$ = 42).

UC, unchallenged; CI, clean injury. Statistical tests: one-way ANOVA with Tukey's post hoc test (B), log-rank test (E, F).

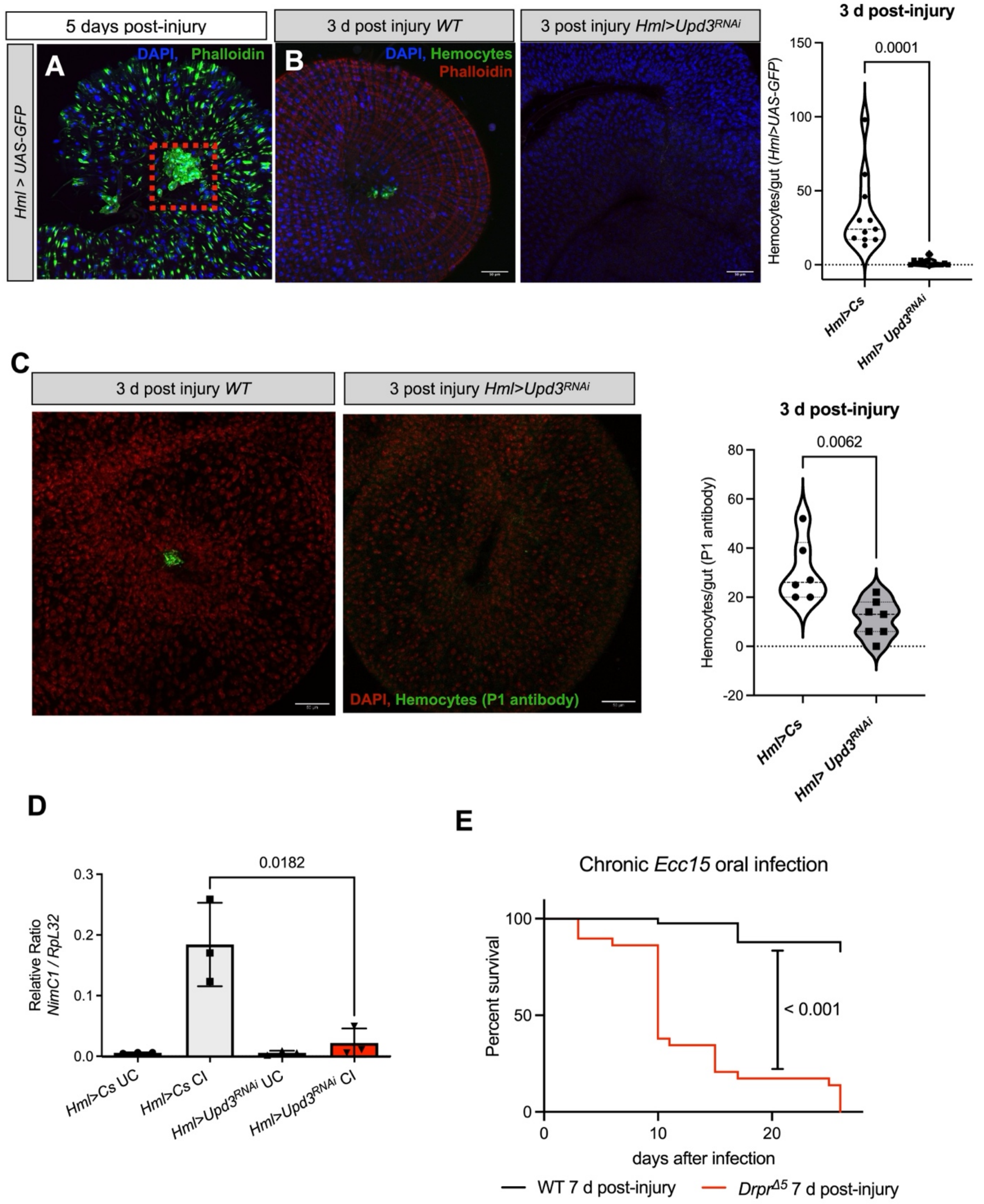


**S5 Fig. Homing of hemocytes to the gut is important to resist an oral infection.**

(**A**) *hmlΔGAL4 >UAS-GFP* fly intestines were dissected 5 days post-injury. The homing of hemocytes was seen to the gut to the loop of R3-R4 regions.

(**B**) Reduced number or even absence of hemocytes are observed in intestines of *hmlΔGAL4 > UAS-Upd3*-RNAi. Quantification of hemocytes that are associated with entire midgut 3 days post-injury from fly intestines of *hmlΔGAL4 > UAS-Upd3*-RNAi flies.

(**C**) P1 immunostaining of unstretched intestines from *hmlΔ-GAL4 > UAS-GFP* female flies 72 h post-injury reveals robust hemocyte recruitment. In contrast, *hmlΔ-GAL4 > UAS-Upd3-RNAi* flies show markedly reduced $P1^+$ hemocyte numbers. *P*-value determined by Student's *t*-test.

(**D**) RT-qPCR from fly intestines of *hmlΔGAL4 > UAS-Upd3*-RNAi flies shows a reduced expression of the hemocyte marker *NimC1* in the adult flies intestines in response to injury, indicating a defect in hemocyte homing. (**E**) Survival of flies following oral infection with *Erwinia caratovora caratovora 15*, seven days post-injury, as indicated in the scheme above in wild-type and $drpr^{\Delta 5}$ mutant animals, shows that the $drpr^{\Delta 5}$mutant is more susceptible. Flies per condition are pooled from at least three independent experiments. Log-rank test used for comparing wild type ($w^{1118}$, n = 50 & $Drpr^{\Delta 5}$, n = 51).
